# Canopy Volume as a Tool for Early Detection of Plant Drought and Fertilization Stress: Banana plant fine-phenotype

**DOI:** 10.1101/2021.03.04.433868

**Authors:** Anna Brook, Yasmin Tal, Oshry Markovich, Nataliya Rybnikova

## Abstract

Irrigation and fertilization stress in plants are limitations for securing global food production. Sustainable agriculture is at the heart of global goals because threats of a rapidly growing population and climate changes are affecting agricultural productivity. Plant phenotyping is defined as evaluating plant traits. Traditionally, this measurement is performed manually but with advanced technology and analysis, these traits can be observed automatically and nondestructively. A high correlation between plant traits, growth, biomass, and final yield has been found. From the early stages of plant development, lack of irrigation and fertilization directly influence developing stages, thus the final crop yield is significantly reduced. In order to evaluate drought and fertilization stress, plant height, as a morphological trait, is the most common one used in precision-agriculture research. The present study shows that three-dimension volumetric approaches are more representative markers for alerting growers to the early stages of stress in young banana plants’ for fine-scale phenotyping. This research demonstrates two different group conditions: 1) Normal conditions; and 2) zero irrigation and zero fertilization. The statistical analysis results show a successfully distinguished early stress with the volumetric traits providing new insights on identifying the key phenotypes and growth stages influenced by drought stress.

## 1. Introduction

Sustainable Development Goals (SDG) is a collection of global goals to attain a better future. Sustainable agriculture is at the heart of this agenda. This goal is responsible for ensuring food production systems and implementing resilient agricultural methods, influencing the increase in production and productivity, assisting in maintaining ecosystems, adjusting to climate change and extreme weather. Thus, simultaneously taking into consideration improvement of land and soil quality. One of the ways that can help achieve sustainable agriculture is with Precision Agriculture (PA), a method to accurately apply the right treatment, in the right place, at the right time.

PA embraces a set of technologies that includes sensors, information systems, machinery analysis and Decision Support Systems (DSS) to optimize production by cost-benefit analysis within the agriculture system. Additionally, with the growing variety of precision equipment and applications, farmers and producers have the opportunity to monitor and analyze more data than before, using the most current knowledge in PA (Figure 1).

**Figure 1.**
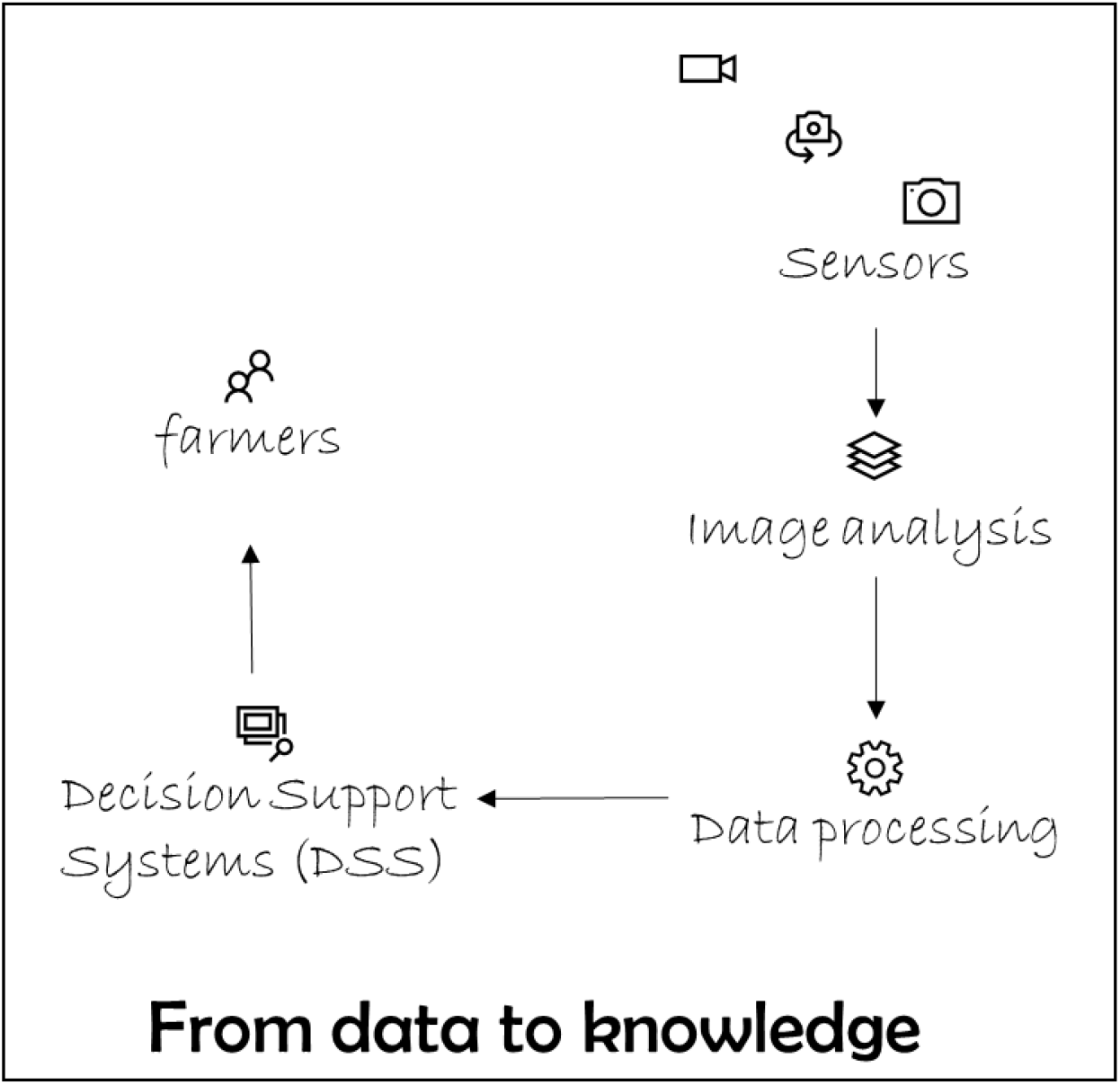
Steps from collecting data for the farmer’s decision.

With the progress in sensors and application in PA, the field of noninvasive plant phenotyping is expanding. Plant phenotyping refers to recording quantitative and qualitative plant traits (Fiorani & Schurr, 2013; Granier & Vile, 2014). To measure plant traits in the traditional manual way is time-consuming, laborious; introducing errors and sometimes destructive (Neilson et al., 2015; Preuksakarn et al., 2010). From 2000s, the era of non-destructive plant-phenotyping platforms, based on image-based techniques, has begun (Granier & Vile, 2014).

Early detection of plant stress are critical to reduce as much as possible productivity loss. Since stress expressed in the leaf surface and structure (Fernández et al., 2008, Shepherd and Wynne Griffiths, 2006) it’s modify plant canopies creating physiological changes, which can be observed and quantify by sensors (Richardson et al., 2013).

Thus, by recording with remote sensing approach, greater opportunities become available for high-throughput trait acquisition and data processing (Dhondt et al., 2013). From as recently as 2010, non-invasive plant phenotyping has significantly progressed with regard to sensors such as fluorescence, visible light, VIS-NIR, thermal, stereo, and tomographic. All contributing to enabling rapid, accurate and non-destructive measurements of plant traits (Qiu et al., 2018; Sankaran et al., 2015; Zhang & Zhang, 2018).

Developing technologies for high-throughput plant architecture capture exposed the need for automated phenotyping methods (Furbank and Tester, 2011). These methods are first extracting plant features and further quantifying biomass and yield (Mathan et al., 2016), assessing drought tolerance (Minervini et al., 2016) and studying morphological properties of plant architectures (Conn et al., 2017; Bucksch et al., 2017; Li et al., 2018). Common phenotyping features among others are size/shape of leaves (Huang et al., 2018) and plant height (Madec et al., 2017; Ziamtsov and Navlakha, 2020).

Plant traits can determine plant performance in terms of growth, final yield, biomass, and adaptation to stress (Dhondt et al., 2013; Gibbs et al., 2019). It is widely considered that estimating above-ground biomass helps in monitoring crop vitality and predicting yield (Bendig et al., 2014; Boukerrou & Rasmusson, 1990; Fischer, 1993). However, since above-ground biomass its extremely difficult to determine non-destructively, other plant parameters are used for this estimation (Tilly et al., 2015). There are many experimental studies, which maintain that plant height is an important morphological trait (Maimaitijiang et al., 2019). This is highly correlated with plant biomass and final yield (*inter alia*, see Table 1 below) (Ehlert et al., 2009; Madec et al., 2017; Qiu et al., 2018). Thus, for demonstrating crop growth, biomass, and final yield, the development of plants over a crop’s life cycle is achieved by inspecting and evaluating morphological traits (Sun et al., 2018).

**Table 1.** Visual and measurable symptoms to water stress in banana crop.

| Stress | Early | Progress |
| --- | --- | --- |
| Leaf-level | Involving a reduction in leaf emergence rate | Yellowing and drying of young leaves |
| Plant level | Reduction in plant height | Pseudostem diameter |
| Bunch level | Delay in bunch initiation | Reduction in yield components |

Light intensity plays a critical role in plant growth and shape-structure changes (Paiva et al., 2003, Hao et al., 2020). As abiotic stress, weak or high light stress both have pivotal effects on crop physiology and morphology. Large and growing studies have investigated that plant leaves usually exhibit particular apparent morphology under different levels of light stress. The study has presented that leaves grown under high light intensity tend to develop thicker than those under low light intensity (Evans and Poorter, 2001, Fu et al., 2012). From Kleinhenz et al. (2003), lettuce is very responsive to light intensity. Distinctive light intensities have an essential effect on chlorophyll and anthocyanins, which lead to visual differences in lettuce. In the experiment by Marschner et al. (1989), chlorosis and necrosis of the leaves were aroused along with the increased light stress. These suggest that it is feasible to grade the light intensity stress of crops by visual analysis.

Sensors and other precision applications have been developed in recent decades with increasing accuracy and are pivotal for estimating plant traits. The near-surface remote sensing technology provides new opportunity for non-destructive, high-efficiency and high-temporal, spatial and in some cases spectral resolution fine phenotyping. In the two-dimension (2D) world, measurements often focus on traits that are not expressed by geometrical measurements. The common plant-phenotyping traits are used to focus mainly on plant structure and color changes via RGB cameras. These types of sensors show progress by adapting new technology independently to the plant-phenotyping world. Multispectral or hyperspectral sensors are mainly used to perform early detection biotic or abiotic stress by radiation from the non-visible spectrum. These sensors collect plant reflectance, which the naked eye cannot be relied upon. Accordingly, hyperspectral sensors cover the spectrum range from Visible and Near-Infrared (VIS–NIR) to Short Wave Infrared (SWIR) in narrow bands. While the multispectral sensors cover specific selected wavelengths in wider bands. Thermal images also acquired from passive radiation, the output data in this case, indicate differentials in temperature from single-leaf to full-plant level, based on the sensor’s spatial resolution (Dhondt et al., 2013; Li et al., 2014; Pieruschka and Schurr, 2019). However, these approaches depend on camera calibration, image registration, and object occlusion. Several platform-specific solutions have been proposed (Nguyen et al., 2016; Gibbs et al., 2018), although general-purpose solutions are absent.

In the three-dimension (3D) world, the measurements provide a more comprehensive and detailed view of the entire architecture and indicate morphological traits (e.g., plant height, number of leaves, volume). Sensors, especially laser scanners, are reliable for extracting morphological traits. One of the great advantages 3D measurements is the generated point cloud, which represents the plant at a certain point in time. There is a variety of information that can be extracted from the point cloud, and not just rely on a single measurement as traditional phenotyping method does. Although 3D devices provide high-resolution and detailed reconstruction, they are only able to measure visible objects but not leaves or fruit concealing one another. This becomes an issue with growing plants and their complex structure starting to overlap. Those problems change from crop to crop and from plant to plant (Quan et al., 2006; Hosoi & Omasa, 2012; Dhondt et al., 2013; Taheriazad et al., 2016 Paulus, 2019; Maimaitijiang et al., 2019). Therefore, developing methods to efficiently process large volumes of point cloud data remains a challenge for 3D plant phenotyping applications as often it affected by non-uniform sampling of points, outliers and missing data and scalability (Chaudhury et al., 2018). Thus, emerging robust computational methods to study plant shapes from 3D point cloud data has become essential.

A 3D visualization of crops for farmers and producers is more intuitive and is a more real-crop observation than the 2D world (Hosoi & Omasa, 2012; Lou, Liu, Han, et al., 2014; Lou, Liu, Sheng, et al., 2014). This way, when the crop is modeled in 3D, it can fill the gaps in farmers and producer’s knowledge and interpret it for the world of automation and machine learning. However, the difficulties encountered in coping with a huge amount of data and large-scale memory resources for analyzing and producing results from the 3D, and spatially in plants where the phenotyping process is occurring, are a new challenge (Nguyen et al., 2016).

Bananas plants (genus *Musa*) is considered an important food security crop with the highest global fruit crop production (130 million tons with agricultural commodity value ~64 billion US$), providing a cheap produced source of energy, rich in certain minerals and in vitamins, and rank as one of the world’s principal consumed food (FAOSTAT, 2018). It is a tropical plant that prefers appropriate temperature and frequent rounds of irrigation. Its production restraints by biotic and abiotic stresses (e.g. drought, salinity and heat). Banana plants are very sensitive to soil water deficit (Robinson, 1996) as its leaves remain highly hydrated even under drought (Turner and Thomas, 1998, Turner, 2004). Therefore, many remote sensing approaches, from image processing to spectral leaf-level analysis, might be less informative for early stress detection.

Nevertheless, according to numerus studies drought is one of the important abiotic constraints restricting banana cultivation and its further adoption into non-conventional growing areas (Ravi et al, 2013). Bananas have a high demand for water with supplemental irrigation often needed to maximize yield (Kumaran and Muthuvel, 2009; Rajak et al., 2017). Considering that water limitation is a major problem for global agriculture affecting almost half of all soils worldwide, it is defined as “shortage of water in the root zone, resulting in decreased crop yield” (Salekdeh et al., 2009). The effect of drought on plants is complex and plants respond (e.g. dehydration and overheating of its cells and tissues) with various protective adaptations.

Early-stress detection and rapid assessment of plant stress symptoms remains acute, especially under drought-stress conditions, which are one of the most important limitations to food production worldwide (Berger et al., 2010). Several recent studies suggested that laser scanner is a supreme tool for nondestructive crop growth monitoring in field and greenhouse practices. Yet, to the best of our knowledge, no study has been conducted to explore the responses of 3D banana plant phenotypes to drought stress using point cloud data. The feasibility of point cloud data in monitoring banana plant phenotype dynamics and how it phenotypes respond to drought stress still need to be evaluated and analyzed. Thus, the object of this study is to focus on morphological characteristics of plants using two groups (control—recommended irrigation and fertilization *versus* zero irrigation and fertilization treatment).

The main aim is to evaluate the performance of 3D data in monitoring time-series banana plants phenotypes in greenhouse practice and analyze the growth dynamics under drought stress. Specifically, two questions were addressed: first, how accurate is point cloud data for banana plant phenotype extraction in greenhouse practice, and how do banana plant phenotypes change under drought stress during the growing period? Second, how can phenotypes associated with early drought stress indicate the occurrence and development of drought in 3D?

## 2. Methodology

Bananas have a high demand for water and high yield will needed a supplemental irrigation (Rajak et al., 2017). Originally grown in tropical regions, cultivation has expanded across a much wider range of both tropical and sub-tropical climates exposing production to extended dry periods (Van Wesemael et al., 2019). Studies showed that a deficit of 100 mm monthly rainfall can reduce bunch weight up to 9% and regions with less than 1100 mm year-1 rainfall might lost between 20–65 % compared to yield potential (Van Asten et al. 2011).

The main banana shoot is a pseudostem structured of the rolled and encircled leaf sheaths that emerge from the compact true stem (Carr, 2009) and growing process of a rapid straight up ejection of a new leaf. Water stress causes oxidative damage and protective mechanisms differ among banana cultivars (Chai et al., 2005). Correlations between stomatal conductance, transpiration, and photosynthesis in water-stressed plants are well documented (Kallarackal et al., 1990). It is visually apparent as reduced growth and reduced leaf size, and increased leaf senescence (Kallarackal et al., 1990; Turner and Thomas, 1998). The rate of emergence of new leaves is the most sensitive indicator of soil water deficit stress (Hoffman and Turner, 1993; Turner and Thomas, 1998). Visual symptoms to water stress are presented in Table 1.

Banana contains laticifers in the leaves, fruit and rhizome and these limits the use of standard methods to measure water relations in the plant (Turner and Thomas, 1998). Several methods for measurements of the thermodynamic leaf water status (Turner et al., 2007) and plant morphological techniques for leaf folding and leaf emergence (Milburn et al., 1990; Turner and Thomas, 1998; Zimmermann et al., 2010) have been developed. However, all the above methods are destructive. Among non-destructive methods for an early water stress detection is a reduction in plant height (Araya et al., 1998). This measure is a well-known in many crops (see Table 2) and very common in agriculture practice as plant height result from drought pressure and directly impacted by abiotic stress and irrigation management.

**Table 2.** Summary of studies reporting significant correlations between plant height and plant condition.

| **Crops** | **Growing conditions** | **Parameter** | **Strength of correlation with**<br>***height*** | **Authors** |
| --- | --- | --- | --- | --- |
| Spring barley | Seven breeding programs | Vegetative biomass | Significantly positive | Boukerrou & Rasmusson, 1990 |
| Barley; oat;<br>Wheat | Five nitrogen-treatments<br>(0, 40, 80, 120 and 160 kg of N/ha) | Grain yield | Significantly positive ($R^2 \approx 0.88-0.95$) | Karjalainen et al., 2008 |
| Rape seed Oil<br>rape<br>Winter rye<br>Winter wheat<br>Grassland | Field conditions | Fresh biomass<br>Dry biomass | Significantly positive<br>($R^2 \approx 0.75-0.99$) | Ehlert et al., 2009 |
| Corn | Six nitrogen-treatments<br>(0, 62, 123, 185, 247, and 308 kg N/ha);<br>no irrigated corn after corn;<br>no irrigated corn after soybean;<br>no irrigated corn after cotton;<br>irrigated corn after soybean | Yield | Significantly positive at 10- and 12-leaf stages | Yin, McClure, Jaja, Tyler, & Hayes, 2011 |
| Barley | Two nitrogen- treatments<br>(40 and 80 kg N/ha) | Fresh biomass<br>Dry biomass | Significantly positive<br>($R^2 \approx 0.81-0.82$) | Bendig et al., 2014 |
| Paddy rice | Varying nitrogen-treatments<br>(incl. 0, 70, 100, 130, 160 kg N/ha) | Biomass | Significantly positive<br>($R^2 \approx 0.86$) | N. Tilly et al., 2014 |
| Tropical forest | Natural conditions | Above-ground<br>Biomass | Significantly positive<br>($R^2 > 0.79$) | Ota et al., 2015 |
| Barley | Two nitrogen- treatments<br>(40 and 80 kg N/ha) | Above-ground<br>biomass | Significantly positive<br>($R^2 \approx 0.85$) | Tilly, Aasen, &<br>Bareth, 2015 |
| Wheat | Irrigated/ Water stress | Above-ground<br>Biomass | Significantly positive across<br>growth stages ($R^2 \approx 0.9$)<br>Intermediate at the flowering stage<br>($R^2 \approx 0.5$) | Madec et al., 2017 |
| | | Yield | Intermediate<br>($R^2 \approx 0.5$) | |

Drought directly influences the flowering stage; the most important step in the fruit-growing process. Drought is also expressed at the leaf edges; as a bending of the leaf edge. Lower fruit yield and diminished fruit shell quality is a result of insufficient irrigation and fertilization.

### 2.1. Greenhouse Experiment and Data Collection

Banana plantlets (Musa Acuminata variety GAL) were grown under normal greenhouse conditions in northern Israel by Rahan-Meristem. The plantlets were exposed to stress and were phenotype for drought tolerant traits.

Two experiments were conducted for developing and testing the suggested methodology. The first experiment was carried out in a net greenhouse at the Rahan Meristem, Ltd. site (33° 5′ 35.24″ N, 35° 6′ 17.16″ E) during mid-August 2018 and used for developing and train the suggested methodology. The second experiment was carried out at the same place conducted under the same conditions during mid-August 2019 and used for testing and validating the suggested approach.

In the greenhouse experiments, banana plants (*Musa*) were grown in pots under two different conditions: (1): Normal/recommended irrigation and normal fertilization; i.e., *control*; and (2): zero irrigation and zero fertilization; i.e., *treatment*. The first experiment took place over a period of 9 days (from 19/08/2018 to 27/08/2018) and 7 days of treatment (from 17/08/2019 and 23/08/2019). During the first two days, the plants from the two groups were treated identically; i.e., were normally irrigated and fertilized, to provide both groups with equal starting conditions. Weather conditions during the experiment were an average 27.3°C with an average 75.5% humidity. The plants were individually monitored by the agronomist and ground truth of plant height and the projected leaf area (PLA) were manually measured. Important to note that most of the early identification of stress are through manual identification. Agronomists observed the leaf to assess the state of crop stress. Although, human vision is comprehensive and unique, it is subject to individual differences in determining the pattern of crop stress symptoms.

During the experiment, plants in each group (15 plants each) were observed in the morning of each day before irrigation *via* video Canon Kiss X4 DLSR camera. For collecting 3D data, we used the video mode of a Canon Kiss X4 DLSR camera, with a 22.3 x 14.9 mm CMOS sensor, which has a diagonal of 26.82 mm (1.06”), a surface area of 332.27 mm^2^, pixel pitch of 4.29 μm, and pixel area of 18.4 μm^2^ and pixel density of 5.43 MP/cm^2^. The camera was mounted on a tripod in nadir view and placed 1 m above the ground, providing a full-frame view of the plants. Point clouds for each plant for each observation day were constructed from the videos by structure from motion (SfM) algorithm implemented in RealityCapture software.

### 2.2. Data Processing

The approach in this dataset was to generate a high-density point cloud based on the video dataset and to calculate morphological traits on the point cloud output. The dataset was processed using the extracted images from the acquired video with high overlap (approximately 80% overlap of the images) and applying a pan sharpening filter on each frame separately compensating motion and blurriness. This step was performed using Image Processing toolbox in Matlab2019b environment.

The processed images were processed by RealityCapture software, as a typical SfM pipeline for dense point cloud estimation images and run in three main steps. The first step detects feature points in the images using a feature detector; e.g., SIFT (for an overview see Lowe, 1999, 2004), and searches for pairwise matches of the features between the input images. The second step, referred to as bundle adjustment, takes the feature points and the images as input and estimates the Pose (position and orientation); the intrinsic parameters (if not calibrated prior to the capture), and the 3D location of the features, using an optimization scheme where the projection error of the 3D points onto the cameras is iteratively minimized (for an overview, see Wu, Frahm, & Pollefeys, 2011). Finally, the third step takes the sparse point cloud and the recovered cameras as input, and uses the SfM algorithm to grow patches and obtain the dense point cloud. The result is a point cloud representation: a set of ***xyz*** coordinates approximating the plant’s surface (Figure 2) (Alenyà et al., 2011; Omasa et al., 2007; Quan et al., 2006).

**Figure 2.**
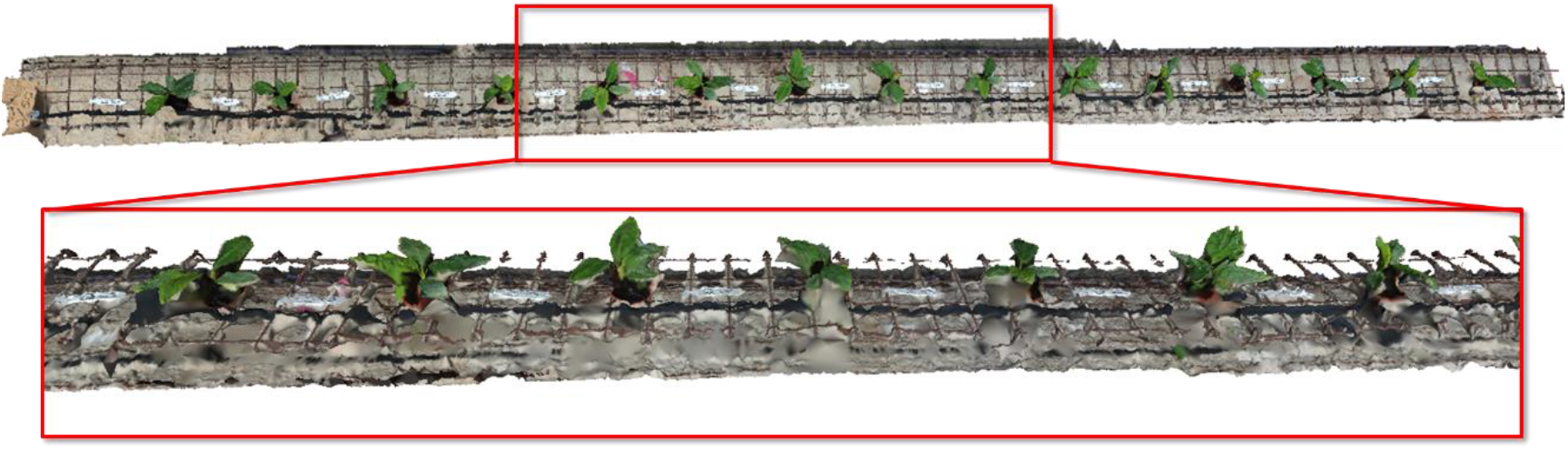
3D model reconstructed from video sequence via Structure from Motion

The resulting point cloud was not arranged on the same coordinate system plane, thus were manually aligned to enable comparing the different dates and treatments used MeshLab software.

As to geo-reference of the aligned point cloud, we extracted the ***xyz*** coordinates of each set of point clouds from the experiment. The background was then cleared and matched by a polynomial function to each point cloud set. At this stage the new ***xyz*** coordinates were determined. This step is performed manually in the Matlab2019b environment. Figure 3, is an example of matching polynomial function.

**Figure 3.**
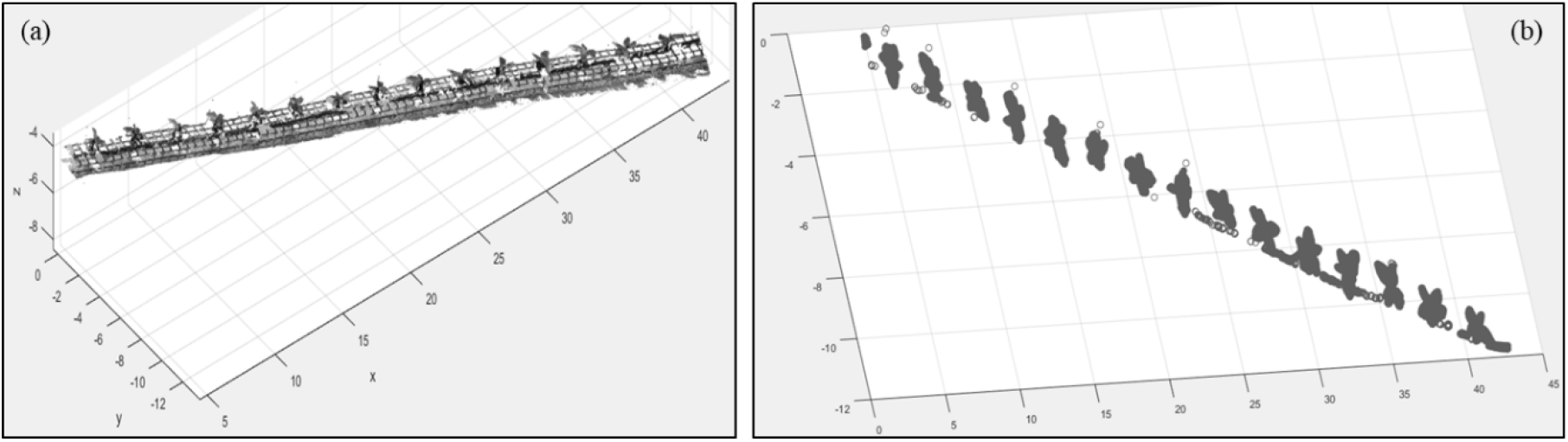
Matching polynomial function; (a) before matching, and (b) after matching.

The last step prior calculating morphological traits, was to normalize point cloud values to validate reference points in the scene, which were manually measured in the greenhouse. Due to the spatial resolution, the different point clouds were non-homogeneous. After normalizing the dataset, each plant was evaluated in the updated spatial resolution by calculating two statistic measurements, i.e. coefficient of determination (R^2^) and root mean-square error (RMSE) between the processed point cloud data and the original point cloud data of validation plants.

To derive morphological parameters for each individual banana plant, it was crucial to first identify and segment each individual plant from point clouds. Because all individual pots were placed in a regular grid with large intervals, by creating a simple grid with a size of 30cm × 30cm and treated the points in each pixel as one banana plant individual. Each plant was separated and the absolute error to determine the differences of the separation for each plant was calculated. Finally, every plant was saved according to date, type of treatment, and number of plants it represented.

### 2.3. Morphological Traits

It has been found that phenotypes related to plant height and leaf area are highly correlated to drought stress (Zhou et al., 2013). Therefore, in this study, the height of each individual plant from the point cloud data of each growth stage for drought stress analysis was calculated as the maximum height from the ground in the corresponding pixel. This parameter is mainly used in 2D and 3D data sets, as it is easy to observe and it should have low deviation errors.

The projected plant area (PLA), which is the projected area of vegetation canopy on the ground was calculating b projecting each individual to the x-y plane, rasterize the projected points and classifying them by the following regulation: pixels with point class 1, and pixels without point class 0. The proportion of class 1 pixels of each banana plant on the x-y plane was the PLA estimation.

The estimation accuracy of plant height and PLA were assessed by the described above coefficient of determination (R^2^) and root mean-square error (RMSE) between the ground truth measurement and the average point cloud-derived estimation of validation plants.

Furthermore, the volume of each plant at each growing period was calculated. Volume is a morphological trait that can help evaluate plant development and health. However, this trait is not possible to evaluate by 2D dataset. Nevertheless, this parameter is mainly used by farmer/agronomist in the field. The following calculation were performed on the point clouds: 1) minimum bounding volume (MVB), 2) standardized linear dimension (SLD), and 3) surface area of minimum bonding volume (SA).

The minimum bounding volume is based on approximations of minimum volume bounding box (MVB) algorithm as shown in Figure 4. This method creates multipath features representing the 3D space. The *concave hull* was selected as a method to determine the fine geometric details of the plant. An example of the result of the mentioned procedure application is reported in Figure 5.

**Figure 4.**
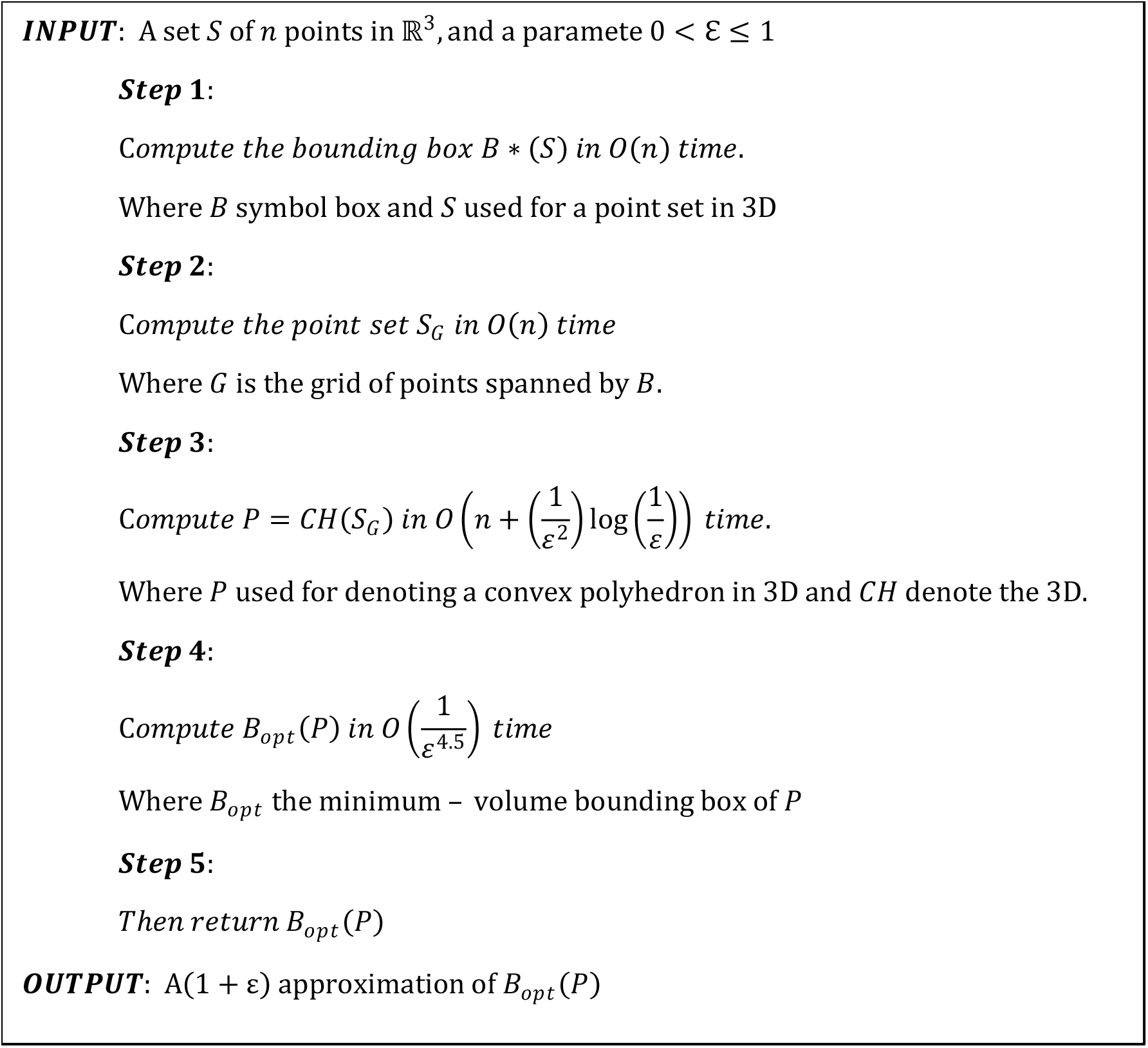
Algorithm approximation of Minimum Volume Bounding box. Source: Barequet & Har-Peled, 2001.

**Figure 5.**
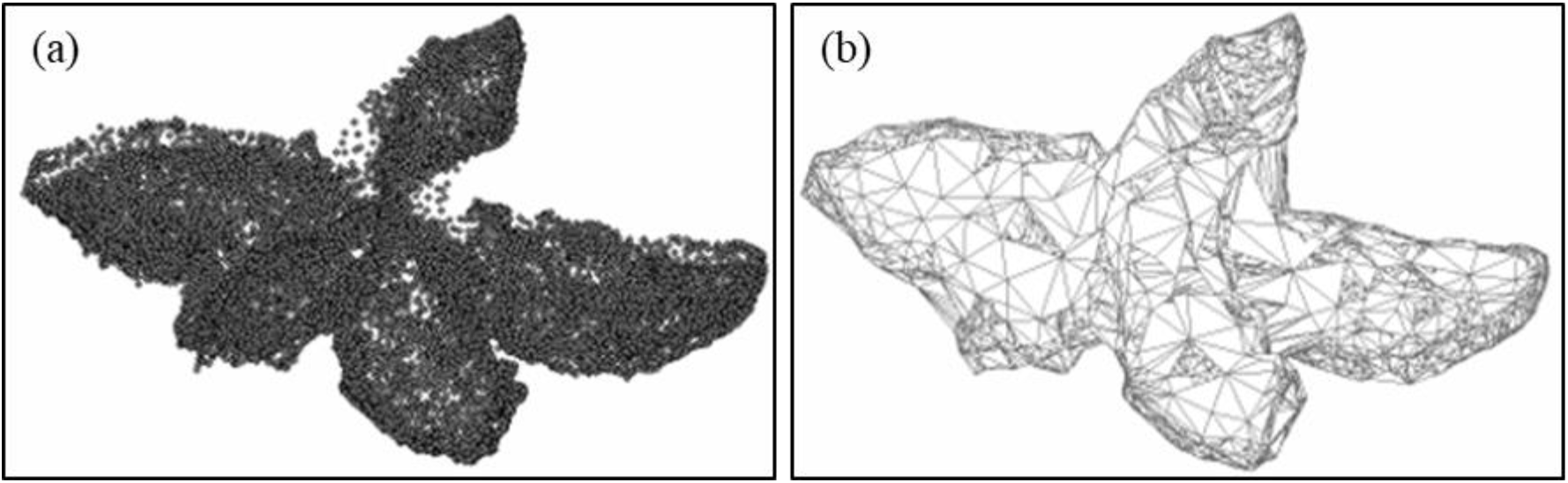
A point-cloud set (a) and a Minimum Bonding Volume (b) of a banana plant at the beginning of the experiment.

Along with PH, PLA and MVB trait, SLD and the surface area of the Minimum Bonding Volume (SA); mining the total area of a 3D surface and standardized linear dimensions, computed as cubic root from the Minimum Bonding Volume, was extracted. It is referred to as the overall dimension of the feature.

The scheme of data collected and point clouds creation, analyze the phenotype dynamics under drought stress, and calculation of PH, PLA, MVB, SLD and SA represent plant height, plant area index, plant area density, minimum bounding volume, standardized linear dimension and surface area of minimum bonding volume for all examined groups (namely *Control and Treatment*) is presented in Figure 5.

**Figure 5.**
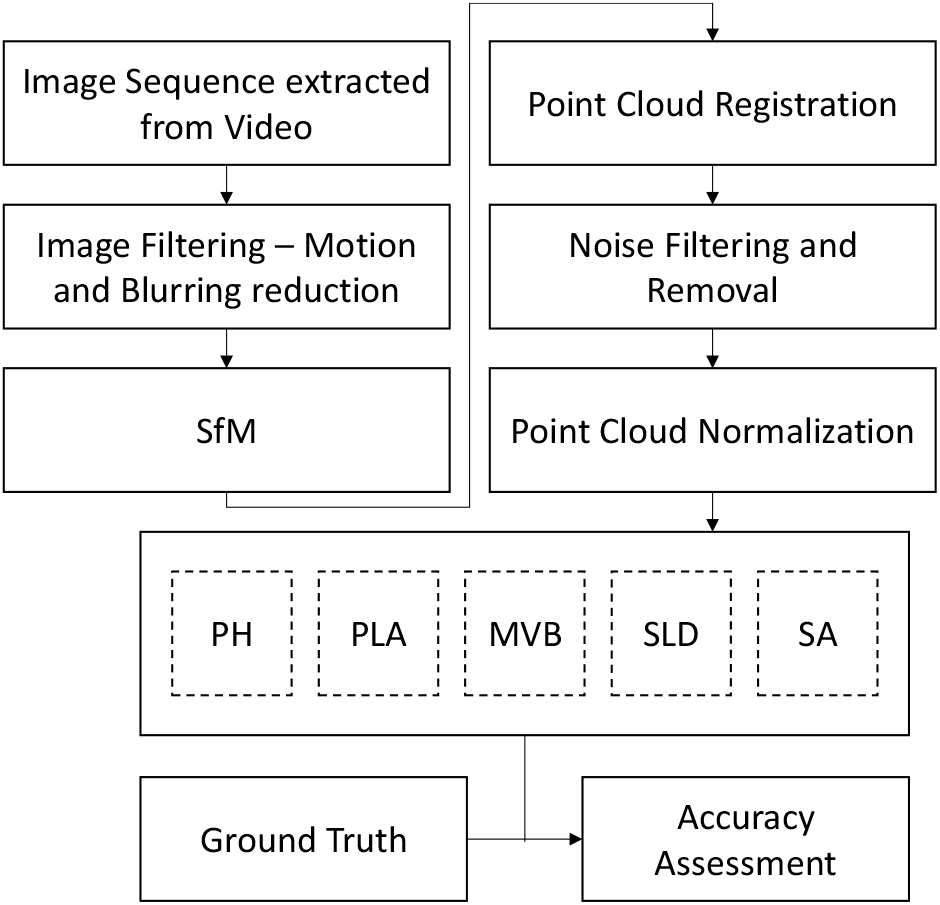
The scheme of data collection, pre-processing and analysis

### 2.4. Analysis of banana plants phenotype dynamics under drought and fertilization stresses

PH, PLA, MVB, SLD, SA measures and the corresponding standard deviations of plants within the same examination group (Control vs. Treatment) were calculated at each growth stage, and the change rates of each parameter compared to the previous stage. The independent sample *t*-tests for the levels of alternative phenotyping traits during seven days of the treatment was calculated and reported. Student’s t-test was used to measure the variability for the control and the treatment groups during the experiment. The null hypothesis was that there was no difference between the values of a phenotype from: 1) two growth groups, 2) between two growth stages. The statistics were used to analyze the change dynamics of phenotypes in different groups. Furthermore, the statistical analysis was implemented to evaluate whether the difference in PH, PLA and three additional traits were significant among growth stages in each group. The time series of MVB, SLD and SA from two studies groups with and without drought treatment were compared to analyze the responses of plant volumetric structures to drought and fertilization stresses.

## 3. Results

The point clouds quality was assessed by two statistical measurements (R^2^ and RMSE) calculated between the processed point cloud data and the original point cloud data of validation plants. The results for 15 individual plants was R^2^ of 0.995 and RMSE of 0.05 cm.

The morphological traits calculated on two banana plant groups; i.e., plants with normal/recommended irrigation and fertilization—*control*, in contrast with plants under stressed irrigation and fertilization conditions over the experiment period; i.e., *treatment*.

The estimated phenotypes were evaluated using measurements of the 30 banana plants collected over eight representative dates (in total 240 samples, equally divided between the control and the treatment groups). Figure 6 shows the conventional phenotypes derived from the 3D data versus measured height and PLA. These results are in good agreements with the ground truth manual measurements. Plant height (PH) had the highest estimation accuracy among the conventional phenotypes calculated based on the processed point clouds results with R^2^ of 0.995 and PL RMSE of 0.217 cm p-value 0.000 for the control group, and with R^2^ of 0.987 and PL RMSE of 0.314 cm p-value 0.000 for the treatment group.

**Figure 6.**
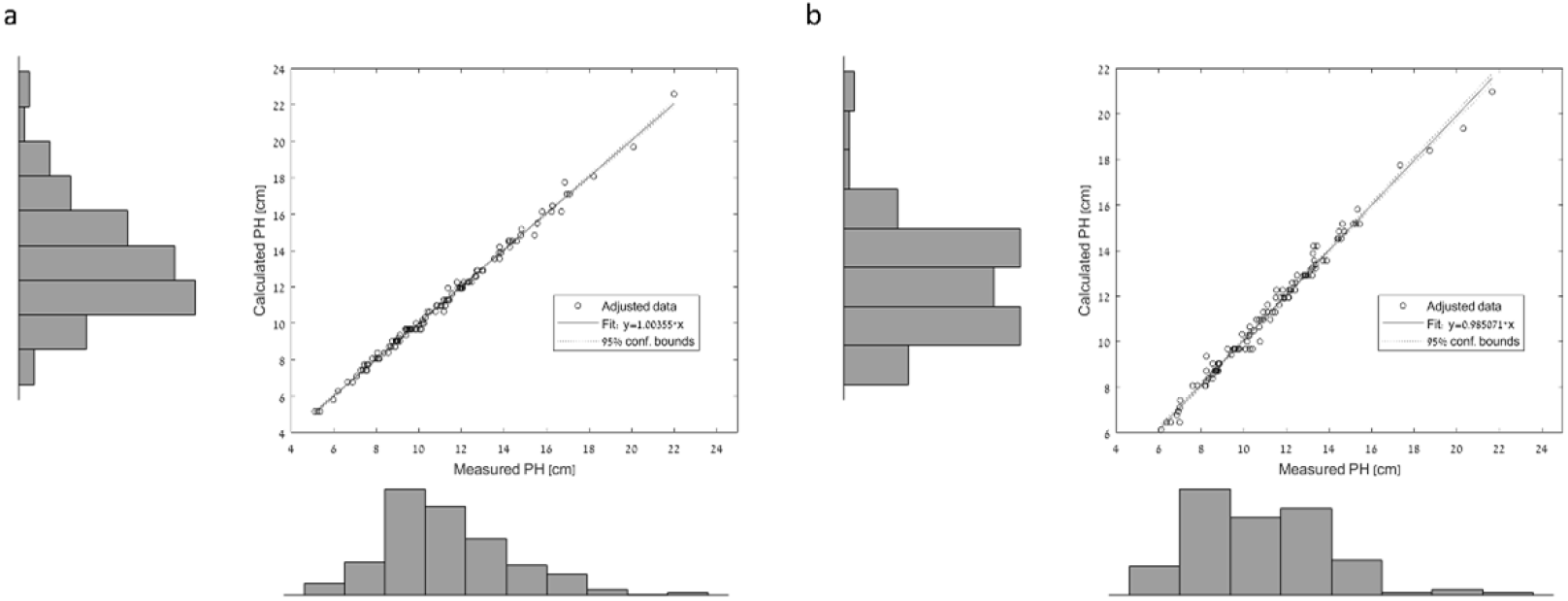
The comparison between the manual measured ground-truth PH and the point cloud-derived estimations for: a. the control group, b. the treatment group.

The PLA estimation results with R^2^ of 0.94 and a RMSE of 0.4 cm^2^/cm^2^ for the control group and R^2^ of 0.92 and a RMSE of 1.5 cm^2^/cm^2^ for the treatment group.

Figure 7 shows the PH in two control days before the experiment. Each plan is numbered and reported, some plants, e.g. 1-3, got lower PH on the second day due to plan general architecture and new leaves, other, e.g. 10-11 grow new leaves on the second day, thus got higher PH values.

**Figure 7.**
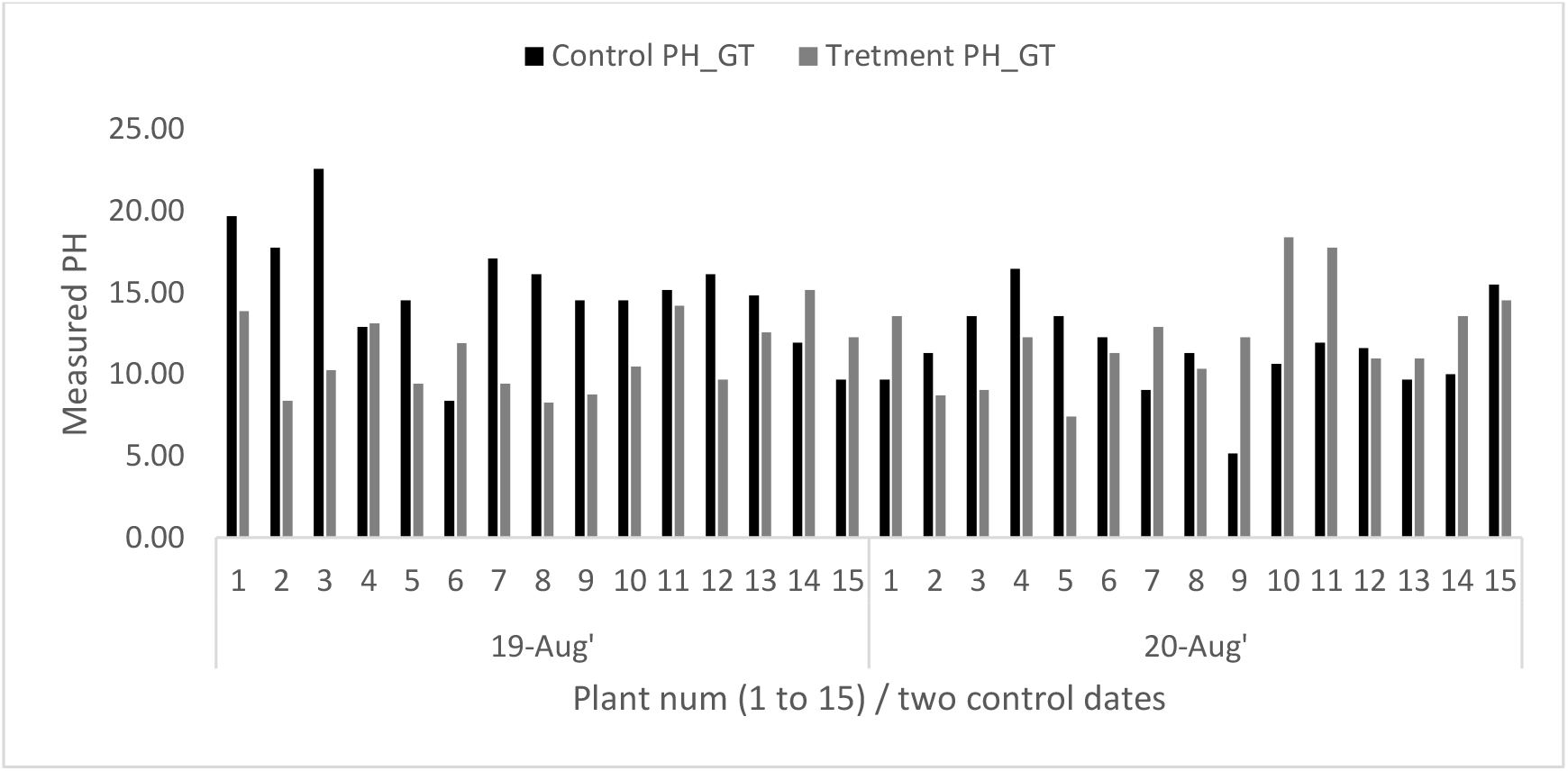
The measured plant height in two control days before the experiment

In average (Fig. 8) the deference between the studied 15 plants was significant (calculated by ANOVA single factor) on the first day (19-Aug’) with p-value 0.012 and became nonsignificant on the second day (20-Aug’) with p-value 0.44 and the experiment could begin.

**Figure 8.**
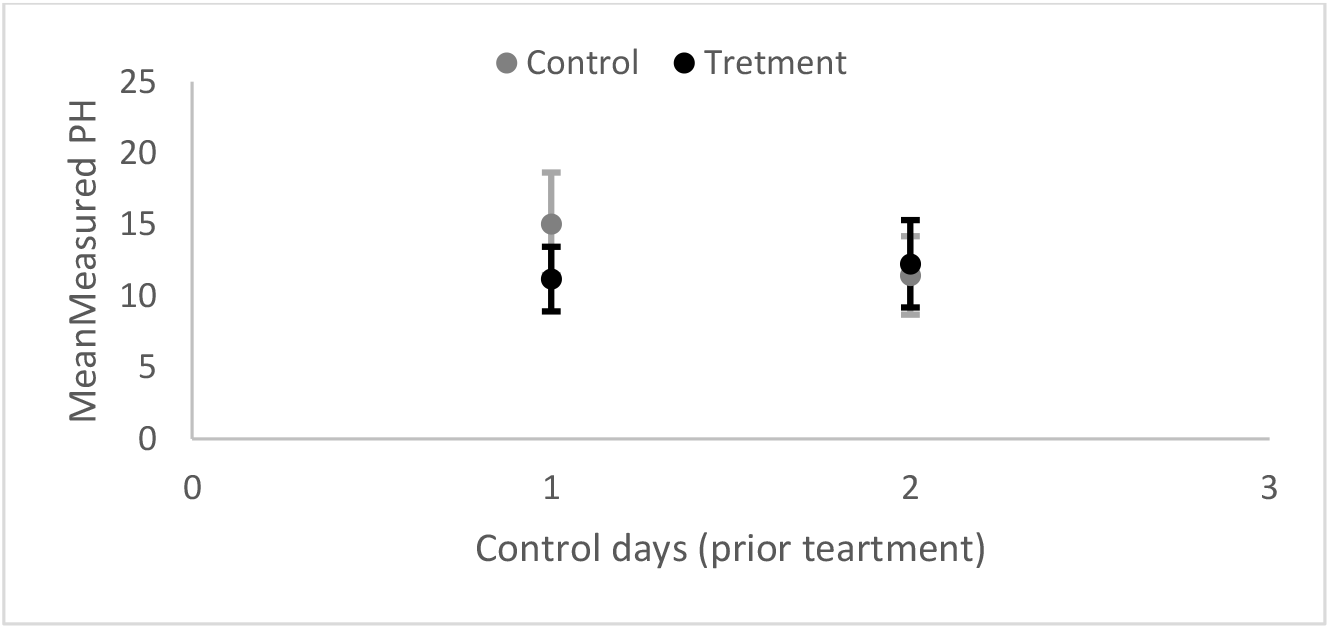
The mean measured plant height in two control days before the experiment

As one can see from Figures 7 and 8, the first two dates of the experiment (19-20/08/2018) were aimed at establishing equal conditions for the plants. It appears that the plants from the control group started at a disadvantage from the treatment group. Similar conditions would be expected on average for the two groups on the five calculated parameters.

During the experiment, seven days of drought and fertilization stress, there was a significant change between the control and the treatment groups (Fig. 9). The control group’s values indicate better growing than the treatment group’s values in four morphological traits (PLA, MVB, SLD and SA). The calculated PH of these groups showed less pronounce difference and the main pattern remains similar between different (control and treatment) groups.

**Figure 9.**
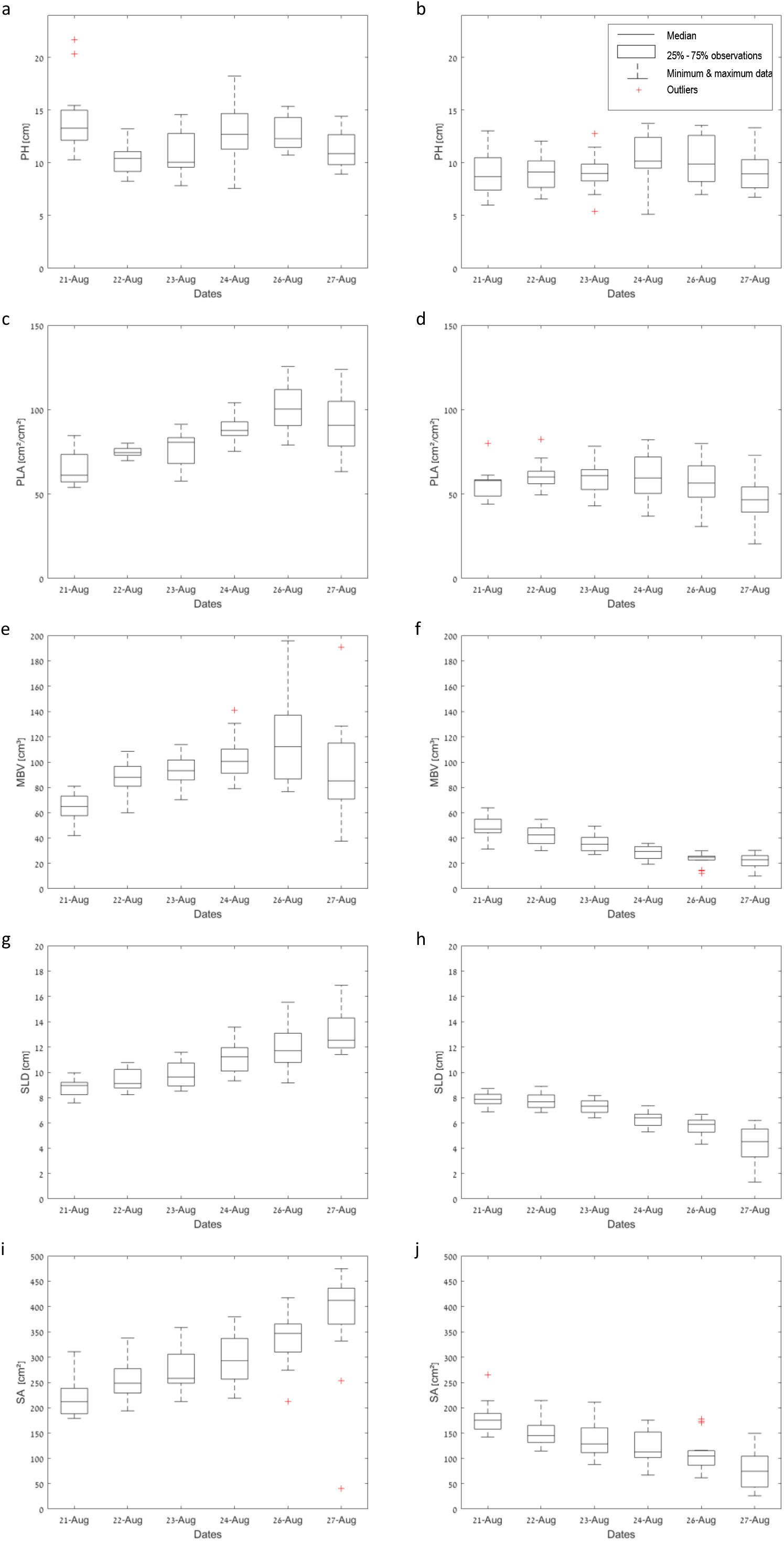
Average dynamics of phenotyping traits of two banana plant classes. Left column represents the control plants and the right one is for the treated ones; **a,b** are display the PH (plant height) in cm; **c,d** the PLA (projected leaf area); **e,f** show the MVB (plant volume); **g,h** SLD (linear dimension) in cm and **i,j** the surface area.

In the control PH was slightly decreasing after the first day, constantly increasing in the following three days, showing no change on the six day slightly decreasing on the last day. Other two calculated phenotypes (PLA and MVB) in the control group followed the pattern of increasing first six days and then slightly decreasing on the last day. The SLD and SA for this group reported continuous growing during the whole experimental period. The PH in the treatment group first three days stayed quite stable showing no change, than slightly increased and immediately decreased. This pattern was similar to the control group. The treatment’s group estimated PLA reported similar pattern on the first three days, indicating no change, and from the fourth day it was showed constant decreasing. The control group had higher PLA growth rate than the treatment group and it kept increasing during the first six days of the experiment. The treatment group’s calculated PLA values stayed lower than the control group during the whole experiment.

In the treatment group the following three phenotypes, namely MVB, SLD and SA, showed the fastest respond to the drought and fertilization stress, and identify abnormal plant behavior at the earlies stages (Fig. 9). Moreover, only those three traits showed continuous decreasing during the whole experimental period, and also reported smallest variation range, by presenting the tightest differentiation between the minimum and the maximum values in the range.

The detailed examination of each plant in the examined groups is reported in Fig. 10 by calculating daily differences for all tested phenotyping traits. According to the reported results (Fig. 10 a1-e1) the PH is not the most illustrative parameter for the examined classes, since its results cannot clearly separate between the control and treated plants. Most of the control plant are losing height between the first and the second day, slowly gain height between the third and the fourth day and lose it again on the seventh day. The treated plants are mostly gaining height on between third and the fourth day and lose it on the last day.

**Figure 10.**
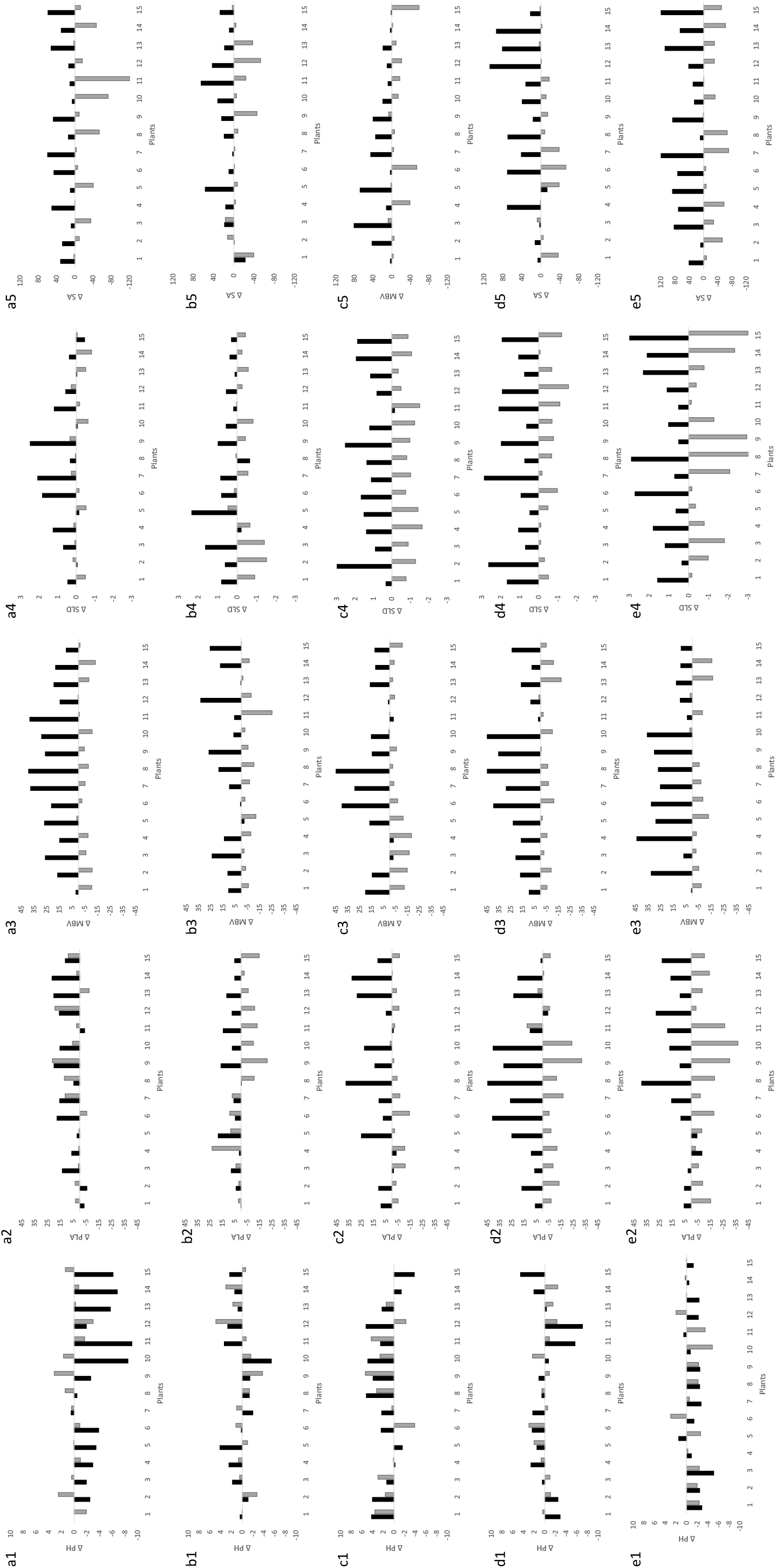
Calculated phenotyping traits differences between two following days for each plant (15 plants in each examined class), in total five pairs. The control plants in black and the treated plants in gray. The **a1-a5** are the PH change (plant height) in cm; **b1-b5** the PLA (projected leaf area) differences; **c1-c5** the MVB (plant volume) variation; **d1-d5** are the SLD (linear dimension) changes in cm and **e1-e5** the surface area differences.

The largest deviation between the control and the treated plants are reported by the SLD measuring the linear dimension of the canopy addressing the changes mostly to leaves. From the third day no control plants showed negative values for this trait and clear separation between classes is illustrated. Furthermore, during the experimental period, as time goes by, the reported deviation become more pronouns as the treated plants values become larger negative and the control plans values become larger positive (Fig. 10 a4-e4).

The next largest deviation between the control and the treated plants are reported by the SA measuring surface area (Fig. 10 a5-e5) and PLA measuring projected leaf area (Fig. 10 a2-e2). Although the PLA shows larger deviance between the groups (negative values in treated plants and positive in control plants) especially during last days of the experiment, the SA trait considered as better evaluator as it shows no confusion between the groups (no control plants with a negative value).

The last is the MVB measuring plant volume is reported in Fig. 10 a3-e3. This trait shows clear deviation between the examined groups, but the values are less distinct. Therefore, according to Fig. 10 the difference between treated and control plants is less noticeable. However, it is important to understand the general concept of this phenotype and its calculation. According to the results reported in Fig. 9, the MVB showed continuous decreasing with the smallest variation range during the experimental period. This observation might explain the results in Fig 10 a3-e3, as the treated plants continuous lose their volume, the calculated difference between two following day’s remains similar. Consequently, the MVB phenotype is presented by similar values during the whole period. In addition it is important to notice that among all the examined traits only two phenotypes showed immediate distinction between the treated and the control plants, namely, MVB and SA, as only these reported clear separation on the second day of the experiment.

Student’s t-test results measuring the variability for the control and the treatment groups during the experiment is reported in Table 3. The PH trait results shows no temporal trends in the examined groups, the p-Value of the treatment group shows insignificant variation during the tested period, meanwhile the control group reports on highly significant difference between plants at the beginning of the experiment. The PLA phenotype reports no variant up to fourth day, and significant differences between the treated plants during three last days of the experiment. The SLD is insignificant only at the first two days, and both MVB and SA traits are significant from the first day of the experiment for the treatment group. The PH, PLA and MVB are trendless for the control group, as its p-Values are partially significant and partially insignificant, without any temporal pattern. However, the SLD and SA traits are insignificant only at the first two days.

**Table 3.**
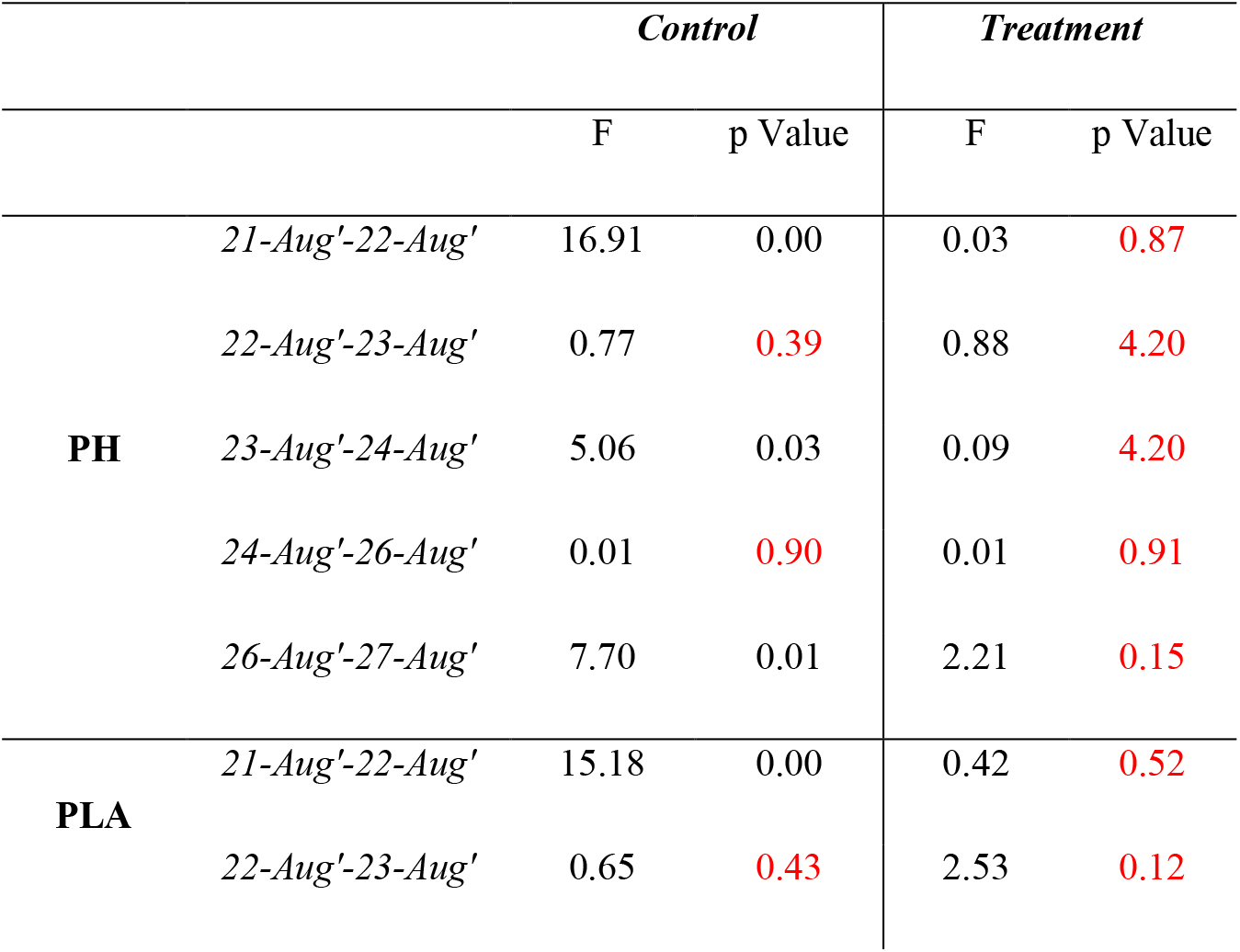
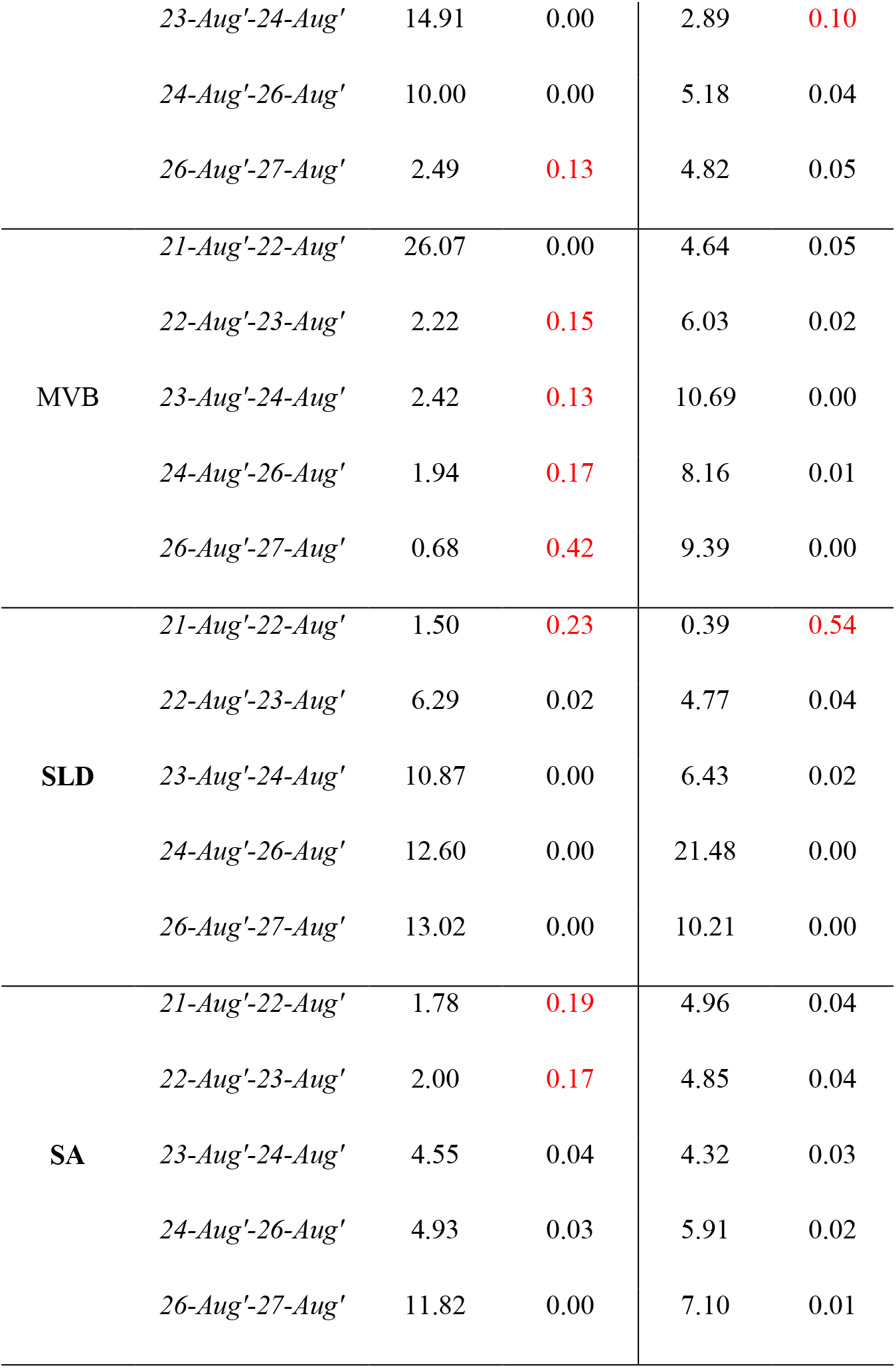
T-test for mean differences between two following days in control and treatment classes separately per trait

Further statistical analysis is implemented between the two examined group on a daily bases for each tested trait. Table 3 summarizes the results of independent sample *t*-tests for the levels of alternative phenotyping traits during the experimental period. Once again Student’s t-test is used to measure the variability between the seven days of the experiment for the control and the treatment groups.

According to the results reported in Table 4, on the first day, significant differences between the two groups in terms of height (PH), and volume (MVB) were reported, while plant dimension (SLD) and plant area (PLA and SA) did not report significant differences. The differences between the control and treatment groups become insignificant on the second, third and the last day of the experiment for the PH and PLA traits, showing that these phenotype cannot distinguish between the examined groups.

**Table 4.**
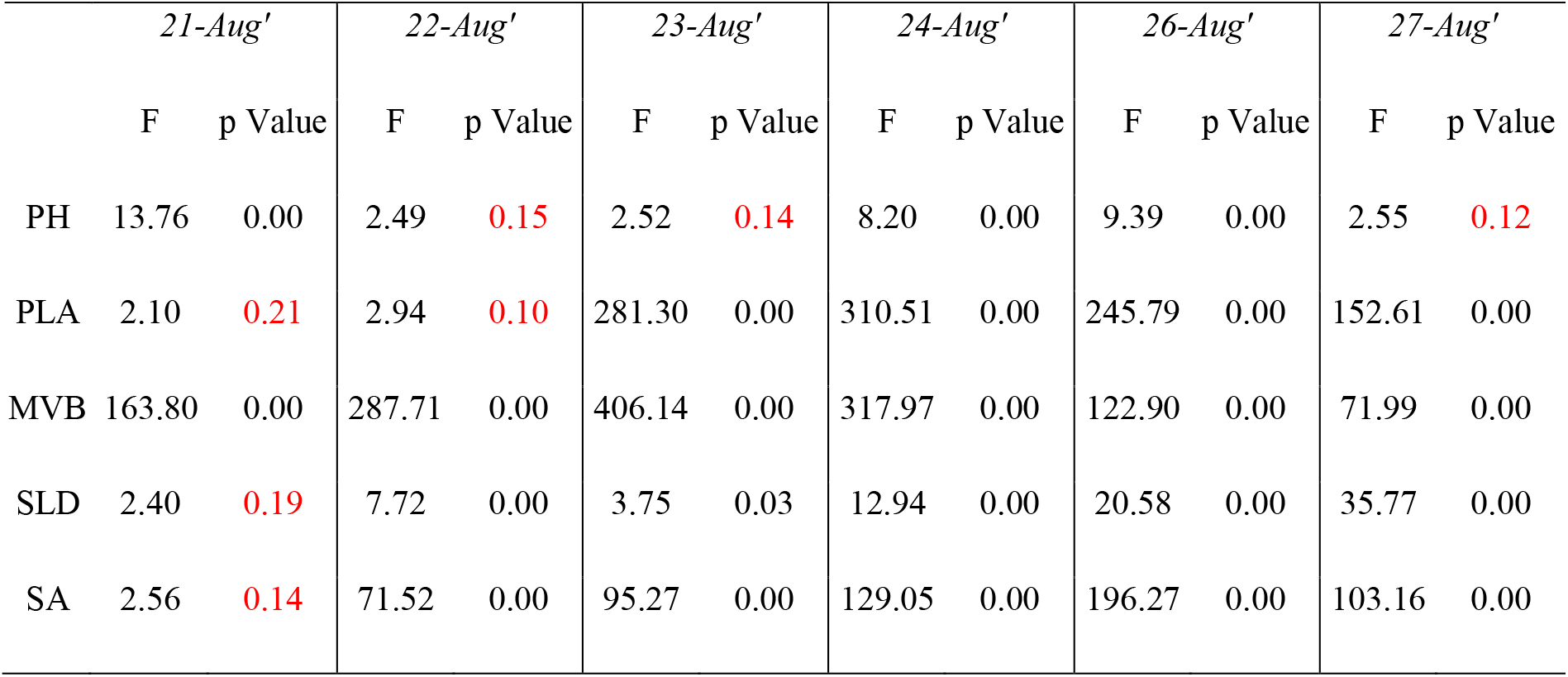
T-test for mean differences between the control and treatment classes per day

The PLA discriminates between the groups only after the second day. However, only MVB separates between control and treatment plants from the first day at a heist significance level (p-Value 0.00).

## 4. Validation and Verification

The second experiment took place at the same greenhouse between 17^th^ August to 23^ed^ August, banana plants (*Musa*) were grown in pots under the two conditions identical to the first experiment: (1): Normal/recommended irrigation and normal fertilization; i.e., *control*; and (2): zero irrigation and zero fertilization; i.e., *treatment*. During the second experiment, plants in each group (15 plants each) were observed in the morning of each day before irrigation *via* RealSense depth camera D435 by Intel. The point clouds were automatically generated by the devise, with no additional processing.

The obtained results confirm the illustrated phenotype reported in section 3. The results presented in figure 11 show the fist and the last day of the experiment, only for PH and MVB traits. In the treatment group the MVB respond to the drought and fertilization stress, and identify abnormal plant behavior by decreasing during growing period. The PH showed similar average phenotype between the first and the last day of the experiment within the treatment group. Additionally, the PH and MVB showed very high growth rate in the control group.

**Figure 11.**
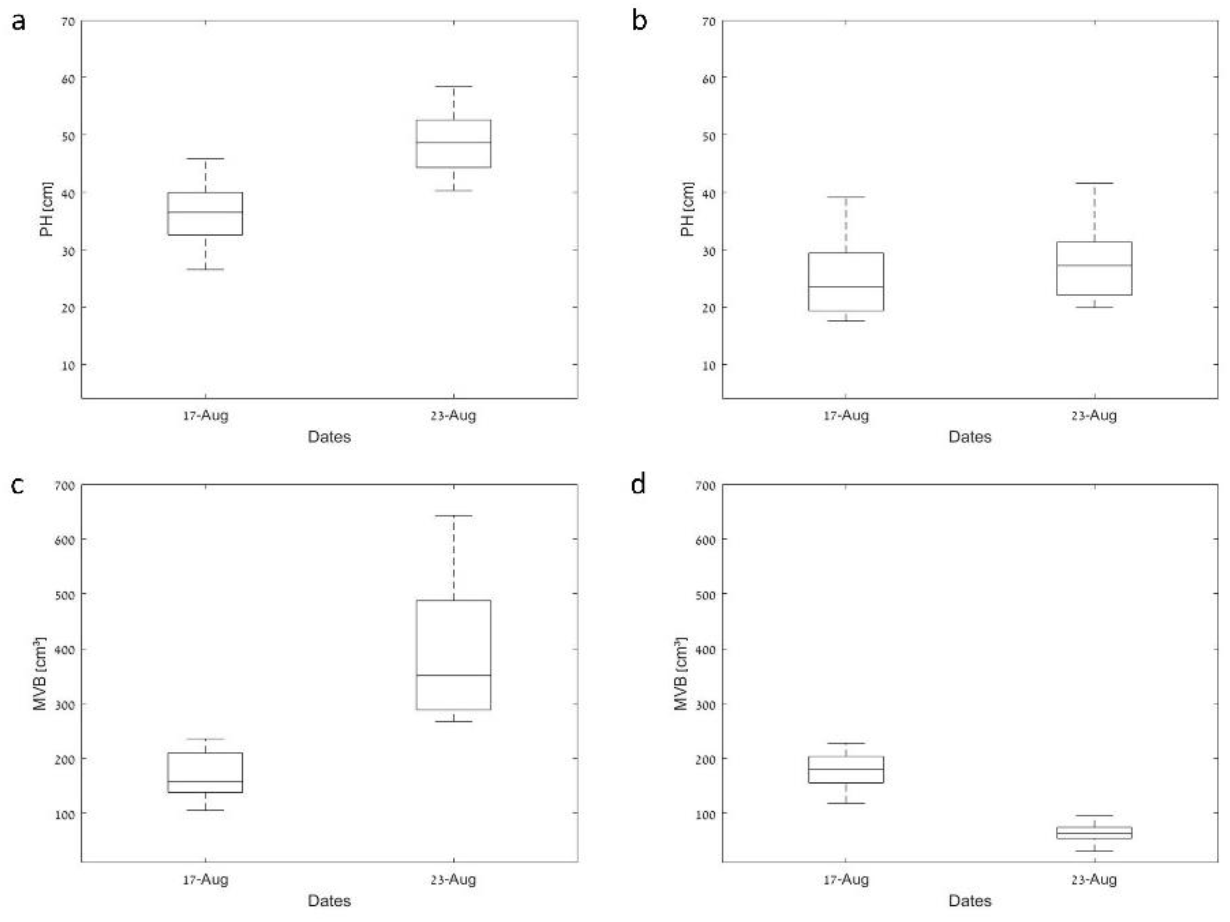
Average dynamics of phenotyping traits of two banana plant classes. Left column represents the control plants and the right one is for the treated ones; **α,b** are display the PH (plant height) in cm; **c,d** the MVB (plant volume).

Table 5 summarizes the results of independent sample *t*-tests for the levels of alternative phenotyping traits during the experimental period. Student’s t-test is used to measure the variability between the first and the last day of the experiment for the control and the treatment groups.

**Table 5.** T-test for mean differences between the control and treatment classes at the first and the last day of the experiment

|  | 17-Aug' |  | 23-Aug' |  |
| --- | --- | --- | --- | --- |
|  | F | p Value | F | p Value |
| PH | 9.88 | 0.00 | 103.09 | 0.00 |
| MVB | 23.98 | 0.00 | 121.42 | 0.00 |

**Table 6.** T-test for mean differences between the first and the last days in the control and the treatment classes separately per trait

|  | *PH* |  | *MVB* |  |
| --- | --- | --- | --- | --- |
|  | F | p Value | F | p Value |
| Control 17-Aug'-23-Aug' | 35.54 | 0.00 | 51.67 | 0.00 |
| Treatment 17-Aug'-23-Aug' | 0.71 | 0.41 | 97.25 | 0.00 |

According to the results reported in Tables 5, on the first day and the last day, significant differences between the two groups in terms of height (PH), and volume (MVB) were reported. However, the examination of each group separately showed insignificant differences within the treatment group for PH trait. The MVB showed significant difference between the groups and within each group between the first and the last days of the second experiment, thus only this trait found to be useful for an early stress detection.

## 5. Discussion and Conclusions

This study investigated and compared two banana plants groups under different conditions: a control group with normal irrigation and fertilization, and a treatment group with no irrigation and no fertilization. The objective: To be able to determine if morphological traits can be used as markers for early detection of drought and fertilization stress.

As the results attest, the levels of such parameters as Minimum Bonding Volume, its surface area, and standardized linear dimension, *do* identify a stress state in its early stage. In particular, the morphological calculation of surface area of Minimum Bonding Volume and its surface area showed the most significant results from the first day of the trial. The proceeding from the scale of the gap in average levels of phenotyping traits, surface area might have comparatively greater potential for stress detection. This probably might be explained because water balance is regulated by evaporation and transpiration processes; both crucially affected by leaf surface area.

The results shows that during the experiment’s short period, in order to apply early detection, the main morphological trait that can lead to detecting stress is volume. This is due to the plant’s manner of growth: extracting new leaves independent of plant stress. When the plant is stressed, the older leaves lose strength at their edges. As a result of the stress, the plant continue growing but the volume is the main traits how affected by that. Therefore, research that focuses mainly on relative changes of average plant height, are incapable of informing of drought and fertilization stress at its early stages, as compared to relative changes in plant volume (Oliveira et al., 2012; Su et al., 2019). In Khanna, et al. (2019) the multi-model datasets of sugar beet (*Beta vulgaris*) under numerous drought stress tests, analyzing several traits, such as height, volume and canopy cover were reported. This paper showed significant changes in height on the 36^th^ day, but it also showed that the volume and canopy cover traits the drought stress was expressed earlier. Nevertheless, there was also a gap between measurements, which were only taken every second week. Thus, as reported in this paper to observe fine-phenotype and slight/minor changes a daily is needed. This emphasizes our position that new research rests upon conventional parameters that take a huge portion of the data analysis in the 2D world; although the available data set can be used to process and analyze data in a multi-modal 3D world.

In the literature, some studies emerge arguing that plant volume is the best representation for growth. In their study, Chen, et al. (2014) tested an Integrated Analysis Platform with near-infrared, visible, and fluorescent images as input data and producing 388 phenotypic traits, including geometric, color-related, FLUO-related, and NIR-related, for monitoring drought stress in barley. They reported that the best marker for morphological traits, and having the highest correlation with manually measured biomass, was digital volume. Therefore, they used volume to model plant growth and viewed it as a representation for above-ground biomass. In another study, Poiré, et al. (2014) investigated nitrogen and phosphorus treatment responses of the model grass, and, as they reported, shoot bio-volume, estimated from the top view and two side views with a RGB camera, the response to available nutrients correlated strongly with bio-volume and growth.

The advantages of point-cloud based calculations were reported for Maize (Su et al., 2019) as a technique to overcome the limitation of phenotyping methods on certain key growth stages, engaging LiDAR (light detection and ranging) sensor in monitoring maize phenotype dynamics at an individual plant level. The results demonstrate the feasibility of using terrestrial LiDAR to monitor 3D maize phenotypes under drought stress in the field.

The way to adopt the most informative parameters that serve the farmers and producers’ goals is not by limit the development of methods and tools to 2-D world. The use of a new approach and technology require a new viewpoint and not fixed only on traits measured in the traditional manner (Njuguna et al., 2010). Therefore, numerous recent papers rise a need for a new traits that can automatically extract high-throughput field-based phenotyping practices from multi-source remote sensing data.

The present study proposed using structure from motion based technology to produce point cloud data and extract temporal banana plant phenotype. Overall, the suggested 3-D traits showed strong capability in assessing a fine-phenotype and early detection of drought and fertilization stress non-destructively and accurately. The study showed that non-destructive and high-accuracy characteristics makes 3-D point cloud based technology an ideal tool in phenotyping applications. The suggested traits may provide new insights on identifying the key phenotypes and growth stages influenced by drought stress.

6.

## Acknowledgment

This research was supported by the Israel Innovation Authority, the PHENOMICS Consortium (grant #67268).

